# Incidence of canine dilated cardiomyopathy, breed and age distributions, and grain-free diet sales in the United States from 2000-2019: A retrospective survey

**DOI:** 10.1101/2020.09.27.315770

**Authors:** Bradley Quest, Stephanie D. Clark, Shiva Garimella, August Konie, Stacey B. Leach, Eva M. Oxford

**Author notes:** Corresponding Author: Eva M. Oxford, DVM, PhD, DACVIM (Cardiology), Chief Research Officer, BSM Partners.

## Abstract

**Background:** Dilated cardiomyopathy (DCM) is considered a predominantly inherited disease in dogs. Recent reports suggest an increased incidence of DCM in atypical breeds eating grain-free and/or legume-rich diets. However, little data regarding incidence of DCM within the US is available; and no existing data quantifies DCM among breeds over time.

**Hypothesis:** We hypothesized that DCM incidence among breeds could be estimated by retrospective polling of veterinary cardiologists. Further, if a correlation existed between grain-free diets and DCM, an increase in DCM would be on trend with increased grain-free pet food sales.

**Materials and Methods:** Thirty-six U.S. cardiology specialty practices were asked for all initial canine and DCM cases evaluated from 2000-2019; fourteen cardiology practices participated. DCM signalment data was provided by three hospitals over 15 years; representing 68 breeds. Age distribution of DCM cases upon diagnosis were compared to other cardio cases and general hospital population from one hospital. All data were evaluated using linear regression models. Grain-free pet food sales data was evaluated from 2011-2019.

**Results:** Fourteen hospitals participated and reported 67,243 unique canine patients. Nationally, data did not support a significant change in percent DCM over time (p=0.85). The overall average incidence rate of DCM during the study period was 3.83% (range 2.41-5.65%), while grain-free diet sales increased 500% from 2011-2019. No correlation between overall DCM incidence and grain free diet sales was discovered. A significant upward trend in mixed breeds diagnosed with DCM, with no significant trend in other breeds was appreciated. An upward trend in age at DCM diagnosis was identified, correlating with trends from overall hospital populations.

**Conclusions:** These data do not support overall increased DCM incidence, or a correlation with grain-free pet food sales. Additional data are necessary to understand whether regional factors contribute to increased DCM within smaller cohorts.

## Introduction

Dilated cardiomyopathy (DCM) is the second most common acquired cardiac disease and the most common cardiomyopathy in the dog and is an important cause of canine morbidity and mortality [1–3]. DCM results in dilation of the cardiac chambers and decreased systolic function, often progressing to congestive heart failure, arrhythmias, and sudden death. While historically considered a predominantly inherited disease common to specific breeds such as the Doberman pinscher [4], Great Dane [5], and Irish wolfhound [6], recent concerns have arisen due to anecdotal reports of an increased number of atypical breeds (those breeds without a known hereditary predisposition) developing DCM phenotype [7–9]. Despite the concerns for increased case numbers, the actual incidence of DCM diagnosed by veterinary cardiologists in the United States is poorly understood, and few historical incidence studies exist [2, 10]. The Veterinary Medical Data Base reported a DCM incidence rate of 0.5% among all dogs from 1986-1991 as reported by veterinary referral hospitals [2]. More recently, a DCM incidence rate of 0.4% was reported among all dogs evaluated at a US veterinary teaching hospital from 1995-2010 [10]. While these studies are helpful to understand overall incidence of DCM, they did not report annual incidence rates of DCM over time.

The question of DCM incidence over time is particularly interesting, given recent reports in the veterinary community of a proposed link between certain diets and DCM. In some cases of DCM diagnoses, the dogs were fed foods that are grain-free and/or high in legumes, which may be produced by small companies [7]. Furthermore, a subset of dogs had improved cardiac function on recheck echocardiograms, after being treated with a combination of medical therapy specific to their disease, taurine and carnitine supplementation, and diet change [8,9]. Additional concerns for an increase in the number of cases of DCM, specifically in atypical breeds, were raised [9]. However, at this time, many questions remain about the role of certain diets in the development of DCM and prospective controlled studies are lacking.

To date, the annual incidence of DCM diagnosed by veterinary cardiologists across the United States has not been reported and no objective data are currently available to evaluate the proportion of different breeds diagnosed with DCM over time. Here, we show the total annual incidence of DCM over the past two decades, as reported by 14 veterinary cardiology services across the United States. Furthermore, our data reveals annual breed distribution of DCM cases from three cardiology services, and annual age distribution from one service. Finally, we report the annual sales of grain-free dog foods over the past decade and discuss the association between grain-free diets and DCM in dogs, as well as alternative hypotheses for the development of DCM phenotype.

## Materials and Methods

### Data Collection

This study represents a retrospective survey of the incidence of DCM diagnosed by veterinary cardiologists from 2000-2019. Thirty-six veterinary referral hospitals with board-certified or residency-trained veterinary cardiologists were contacted to represent all geographic areas throughout the United States. These referral hospitals included veterinary teaching hospitals at accredited colleges of veterinary medicine, as well as private veterinary referral practices. Services were provided with a DCM Incidence Survey (Supplemental Fig 1) and asked to report on how DCM was diagnosed in their facility, how the cases provided were searched through their databases, number of cardiologists at the practice, number of cardiology residents at the practice, search terminology used to identify DCM cases, and criteria used to diagnose DCM.

### DCM incidence

Services were asked to provide the number of initial (unique) canine cardiology cases seen per year; as well as the number of those cases that had an initial diagnosis of DCM. Recheck appointments of the same patient were excluded. Data were presented in clustered bar charts to provide an overall summary. In addition to this data, one hospital provided all annual initial canine cases (all specialties) presenting to the hospital each year, and these data were analyzed.

### Breed distribution

Breed distribution data was provided by three cardiology services (University of Missouri, Columbia, Missouri; Garden State Veterinary Specialists, Tinton Fall, NJ; Red Bank Veterinary Hospitals, Tinton Falls, NJ). A total of 68 breeds were identified from 2004-2019 across the three hospitals. Breeds were divided into six groups: Inherited (breeds with a reported inherited predisposition to DCM) Small breed (< 13.6 kg), Mixed Breeds, Retrievers (Golden Retrievers, Labradors, Chesapeake Bay Retrievers), Other Breeds, and Not Specified. Groups were plotted as stacked plots for ease of visualization, and as scatter plots to evaluate trends over time.

### Age distribution

Age distribution data was provided by Red Bank Veterinary Hospital, Tinton Falls, NJ, from 2005-2020. Data were evaluated by comparing the average age of DCM patients upon initial diagnosis to other canine cardiac patients and other canine patients evaluate by all services. Further, percentage of DCM patients diagnosed at age 0-6 years and 7 + years was evaluated over time.

### Pet food sales data

Growth in annual grain-free pet food sales were provided by The Nielsen Company [11] for the years 2011-2019. Additionally, percent market share of dry dog food was provided from 2016-2019 (The Nielson Company, AXCO, Petco, and Petsmart) [11–13]. These data are shared with permission. We also obtained data concerning the total number of dogs in the US over the past decade [14]. Our goal was to estimate the percentage of dogs eating grain-free food in the US in 2011 and 2019. The percentage of dogs in 2011 was estimated using the following equation: $**900,000,000 grain-free sales / $50.00 per bag / 12 months / 78,000,000 dogs = 2%**. This considered the grain-free pet food sales data available from pet-specialty stores only [11]. This equation was repeated for 2019: $**5,5000,000,000 grain-free sales / $50.00 per bag / 12 months / 90,000,000 dogs = 10%.** Additional data regarding percent of market share of grain free dog food was also considered and resulted in using a range to estimate the percentage of dogs eating grain-free food in 2019. These data are not available for 2011.

### Statistical Analysis

The DCM incidence data were analyzed using Statistical Analysis Software (SAS Institute Inc. 2012. SAS/STAT^®^ 9.4 User’s Guide, Second Edition. Cary, NC: SAS Institute Inc) and expressed as the percent of DCM cases with respect to the number of referrals for each year at each state site. Response was statistically analyzed through a mixed model random coefficient regression analysis where states were designated as the subjects. A linear overall response was assumed across years. The best fitted regression model assumed a common slope (response trend) allowing for differing intercepts among states. States were evaluated together, and in two groups, based on the trends observed. Significance was set at p < 0.05.

Dog breeds were grouped by kind and groups were evaluated using a Poisson regression to analyze breed distributions. The number of dogs in each of the five classes (Inherited, Small Breed, Retrievers, Other, or Mixed Breed) were modelled as a Poisson distributed response variable in a generalized linear model. We modelled the year of each observation as a random effect, controlling for the total number of dogs seen by the cardiology service in each year and the vet hospital (Red Bank, Garden State, or Missouri). Our primary interest was the regression coefficient, its confidence interval, and its p-value. Significance was set at p < 0.05. Computations in the breed analysis were performed using the R statistical package.

## Results

Of the 36 cardiology services approached, 22 declined to participate because of various reasons such as being unable to search their medical record systems, or not having the resources to do such a search. Fourteen services agreed to provide records and were located in the following geographic areas: East Coast (n=6), Pacific Northwest (n=1), Hawaii (n=1), Midwest (n=5), Southeast (n=1) (Fig 1A). Of the 14 cardiology services that provided data, one provided 19 years of data, two provided 15 years of data, four provided between 8-10 years of data, and the remaining seven services provided five years or less of data (Table 1). Cardiologists reported a variety of methods in diagnosing DCM, as listed in the DCM Incidence Survey (Supplemental Fig 1); all reported using breed specific reference intervals for left ventricular internal diameter in diastole (LVIDd) and left ventricular internal diameter in systole (LVIDs), fractional shortening (FS) <25%, and breed predisposition as benchmarks for diagnosis. Other parameters were used less frequently for diagnosis of DCM. All database searches included a diagnosis of “DCM”, or “Dilated Cardiomyopathy”, and no services searched specifically for cases with a diagnosis of “Systolic Dysfunction”. Percentage of total initial DCM cases versus the general hospital population were plotted for one hospital (Red Bank – Tinton Falls, NJ) and correspond with previously published data [2,10] (Fig 1B). An upward trend was noted in both percentage of total DCM cases and percentage of total initial cardiology cases vs the general hospital population during this time period, (Figs 1B - C), indicating that the increase in DCM cases corresponded to an overall increase in cardiology caseload. Annual average percentage of DCM was 3.83% (Range 2.41-5.65%, STD +/- 0.77%) (Fig 1D). A mixed model random coefficient regression analysis revealed no upward or downward trend in DCM incidence when all service sites were considered as a whole (Slope = −0.0066, p=0.8492) (Fig 2). One site, Ethos – Hawaii, provided only one year of data and was excluded from the regression analysis. When sites were considered individually, two different trends were appreciated. These sites were grouped based on their negative or positive trends. Seven services (NC, NY, NJ (n = 2), WI, FL, KS) revealed a negative trend that was statistically significant (Slope −0.1115, p = 0.019). Six services (CO, IL, MA, MO, NH, WA) revealed a positive trend that was not statistically significant (Slope 0.0563, p=0.2703) (Supplemental Table I).

**Figure 1.**
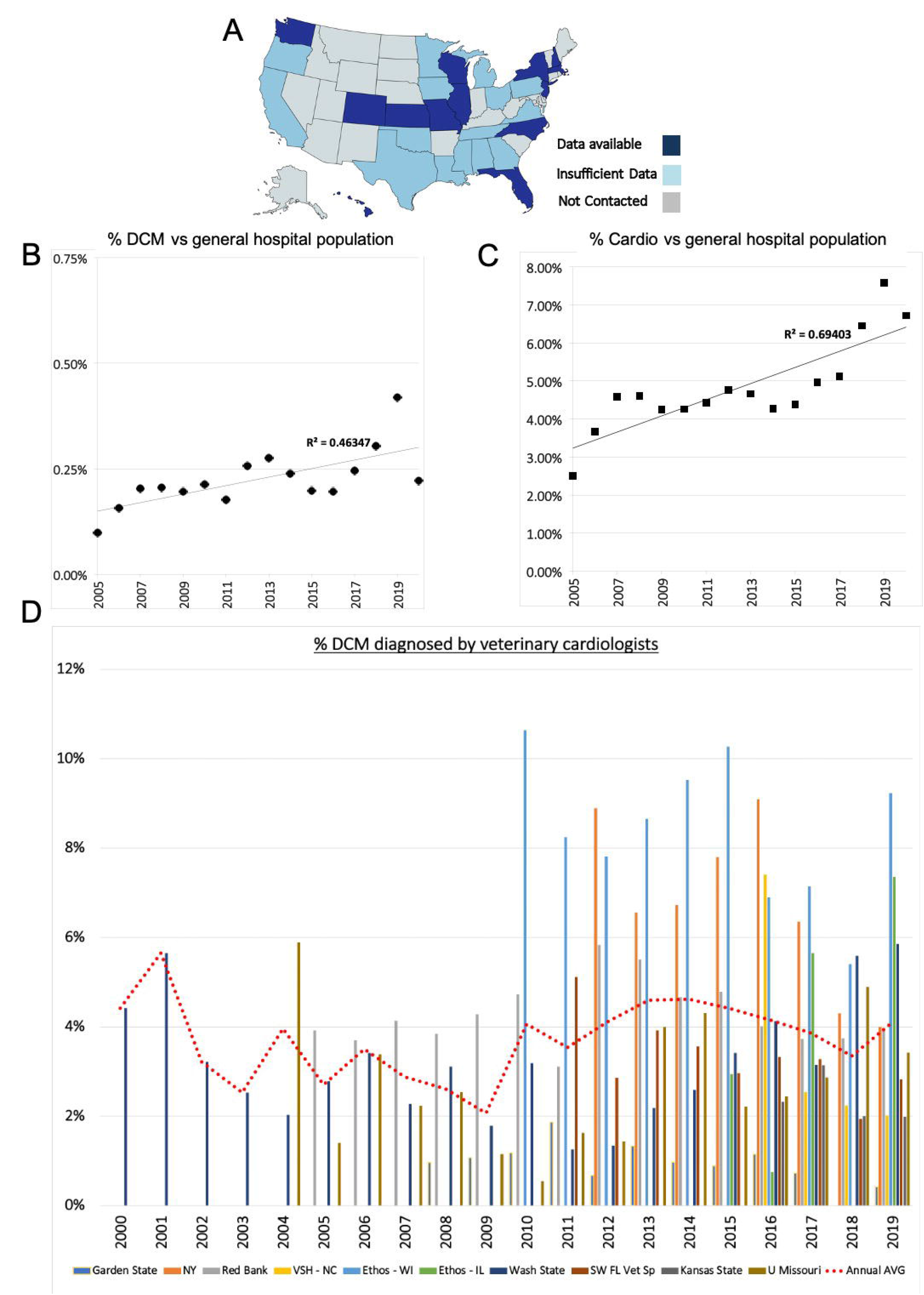
DCM trends in the US. Panel A. Statewide distribution of cardiology services participating in the survey. Light blue represents services that declined to participate. Dark blue indicates the location of participating services. Grey indicates states that did not have a cardiology service contacted. Panels B and C show scatter plots indicating % DCM cases vs the general hospital population (Panel B), and % total cardiology cases vs general hospital cases (Panel C) at Red Bank, Tinton Falls, NJ from 2005-2020. Percentages in Panel B correspond with previously reported estimates of DCM incidence. Linear regression trend lines are displayed Panel B, R=0.46347, Panel C, R=0.69403). Panel C shows that the increase in DCM cases noted over time is a function of the increase in total cardiology cases over the time period. Panel D. Bar chart representing overall DCM incidence reported by 10 cardiology services providing 4 or more years of data. Red dotted line denotes average annual incidence among states. Year on year average for all services was 3.83% (Range 2.41-5.65%, STD + / - 0.77%).

**Table 1.** Annual DCM cases and annual initial cases reported by 14 cardiology services from 2000-2019.

|  | 2000 | 2001 | 2002 | 2003 | 2004 | 2005 | 2006 | 2007 | 2008 | 2009 | 2010 | 2011 | 2012 | 2013 | 2014 | 2015 | 2016 | 2017 | 2018 | 2019 | Total<br>cardiology<br>referral<br>cases |
| --- | --- | --- | --- | --- | --- | --- | --- | --- | --- | --- | --- | --- | --- | --- | --- | --- | --- | --- | --- | --- | --- |
| Cornell/VSC |  |  |  |  |  |  |  |  |  |  |  |  |  |  |  |  |  |  |  |  |  |
| ANNUAL DCM |  |  |  |  |  |  |  |  |  |  |  |  | 37 | 8 | 35 | 39 | 36 | 15 | 26 | 15 | 211 |
| ANNUAL INITIAL CANINE |  |  |  |  |  |  |  |  |  |  |  |  | 416 | 122 | 520 | 500 | 396 | 236 | 604 | 375 | 3169 |
| Ethos-Colorado |  |  |  |  |  |  |  |  |  |  |  |  |  |  |  |  |  |  |  |  |  |
| ANNUAL DCM |  |  |  |  |  |  |  |  |  |  |  |  |  |  |  |  |  | 7 | 33 | 54 | 94 |
| ANNUAL INITIAL CANINE |  |  |  |  |  |  |  |  |  |  |  |  |  |  |  |  |  | 336 | 802 | 1325 | 2463 |
| Ethos-Hawaii |  |  |  |  |  |  |  |  |  |  |  |  |  |  |  |  |  |  |  |  |  |
| ANNUAL DCM |  |  |  |  |  |  |  |  |  |  |  |  |  |  |  |  |  |  |  | 26 | 26 |
| ANNUAL INITIAL CANINE |  |  |  |  |  |  |  |  |  |  |  |  |  |  |  |  |  |  |  | 660 | 660 |
| Ethos-Illinois |  |  |  |  |  |  |  |  |  |  |  |  |  |  |  |  |  |  |  |  |  |
| ANNUAL DCM |  |  |  |  |  |  |  |  |  |  |  |  |  |  |  | 2 | 1 | 20 | – | 32 | 55 |
| ANNUAL INITIAL CANINE |  |  |  |  |  |  |  |  |  |  |  |  |  |  |  | 68 | 134 | 354 | – | 435 | 991 |
| Ethos-Massachusetts |  |  |  |  |  |  |  |  |  |  |  |  |  |  |  |  |  |  |  |  |  |
| ANNUAL DCM |  |  |  |  |  |  |  |  |  |  |  |  |  |  |  |  |  |  | 16 | 26 | 42 |
| ANNUAL INITIAL CANINE |  |  |  |  |  |  |  |  |  |  |  |  |  |  |  |  |  |  | 1298 | 1248 | 2546 |
| Ethos-New Hampshire |  |  |  |  |  |  |  |  |  |  |  |  |  |  |  |  |  |  |  |  |  |
| ANNUAL DCM |  |  |  |  |  |  |  |  |  |  |  |  |  |  |  |  |  | 6 | 46 | 74 | 126 |
| ANNUAL INITIAL CANINE |  |  |  |  |  |  |  |  |  |  |  |  |  |  |  |  |  | 55 | 483 | 510 | 1048 |
| Ethos-Wisconsin |  |  |  |  |  |  |  |  |  |  |  |  |  |  |  |  |  |  |  |  |  |
| ANNUAL DCM |  |  |  |  |  |  |  |  |  |  | 25 | 22 | 20 | 27 | 28 | 31 | 22 | 23 | 14 | 24 | 236 |
| ANNUAL INITIAL CANINE |  |  |  |  |  |  |  |  |  |  | 235 | 267 | 256 | 312 | 294 | 302 | 319 | 322 | 259 | 260 | 2826 |
| Garden State NJ |  |  |  |  |  |  |  |  |  |  |  |  |  |  |  |  |  |  |  |  |  |
| ANNUAL DCM |  |  |  |  |  |  |  |  | 2 | 9 | 12 | 17 | 7 | 15 | 11 | 10 | 13 | 5 | 0 | 2 | 103 |
| ANNUAL INITIAL CANINE |  |  |  |  |  |  |  |  | 208 | 844 | 1021 | 910 | 1044 | 1126 | 1134 | 1127 | 1131 | 686 | 487 | 476 | 10194 |
| Kansas State |  |  |  |  |  |  |  |  |  |  |  |  |  |  |  |  |  |  |  |  |  |
| ANNUAL DCM |  |  |  |  |  |  |  |  |  |  |  |  |  |  |  |  | 13 | 22 | 15 | 18 | 68 |
| ANNUAL INITIAL CANINE |  |  |  |  |  |  |  |  |  |  |  |  |  |  |  |  | 560 | 700 | 750 | 907 | 2917 |
| Red Bank NJ |  |  |  |  |  |  |  |  |  |  |  |  |  |  |  |  |  |  |  |  |  |
| ANNUAL DCM |  |  |  |  |  | 12 | 22 | 41 | 37 | 38 | 38 | 23 | 46 | 42 | 33 | 35 | 35 | 31 | 40 | 50 | 523 |
| ANNUAL INITIAL CANINE |  |  |  |  |  | 306 | 595 | 991 | 963 | 888 | 804 | 740 | 789 | 763 | 708 | 731 | 873 | 831 | 1069 | 1267 | 12318 |
| SW Florida Vet. Specialists |  |  |  |  |  |  |  |  |  |  |  |  |  |  |  |  |  |  |  |  |  |
| ANNUAL DCM |  |  |  |  |  |  |  |  |  |  |  | 24 | 15 | 24 | 22 | 21 | 27 | 32 | 18 | 22 | 205 |
| ANNUAL INITIAL CANINE |  |  |  |  |  |  |  |  |  |  |  | 469 | 525 | 612 | 618 | 709 | 813 | 977 | 929 | 779 | 6431 |
| U. of Missouri |  |  |  |  |  |  |  |  |  |  |  |  |  |  |  |  |  |  |  |  |  |
| ANNUAL DCM |  |  |  |  | 2 | 5 | 16 | 10 | 11 | 5 | 2 | 6 | 6 | 18 | 19 | 12 | 15 | 21 | 41 | 34 | 223 |
| ANNUAL INITIAL CANINE |  |  |  |  | 34 | 356 | 473 | 448 | 434 | 433 | 366 | 369 | 418 | 450 | 441 | 541 | 613 | 732 | 838 | 993 | 7939 |
| VSH North Carolina |  |  |  |  |  |  |  |  |  |  |  |  |  |  |  |  |  |  |  |  |  |
| ANNUAL DCM |  |  |  |  |  |  |  |  |  |  |  |  |  |  |  |  | 4 | 17 | 21 | 20 | 62 |
| ANNUAL INITIAL CANINE |  |  |  |  |  |  |  |  |  |  |  |  |  |  |  |  | 54 | 671 | 939 | 992 | 2656 |
| Washington State |  |  |  |  |  |  |  |  |  |  |  |  |  |  |  |  |  |  |  |  |  |
| ANNUAL DCM | 8 | 14 | 14 | 12 | 13 | 16 | 14 | 10 | 14 | 10 | 19 | 9 | 9 | 14 | 15 | 19 | 23 | 22 | 42 | 53 | 350 |
| ANNUAL INITIAL CANINE | 181 | 248 | 435 | 475 | 640 | 576 | 410 | 440 | 450 | 560 | 596 | 715 | 670 | 640 | 580 | 556 | 557 | 699 | 752 | 905 | 11085 |
|  |  |  |  |  |  |  |  |  |  |  |  |  |  |  |  |  |  |  |  |  | 67243 |

**Figure 2.**
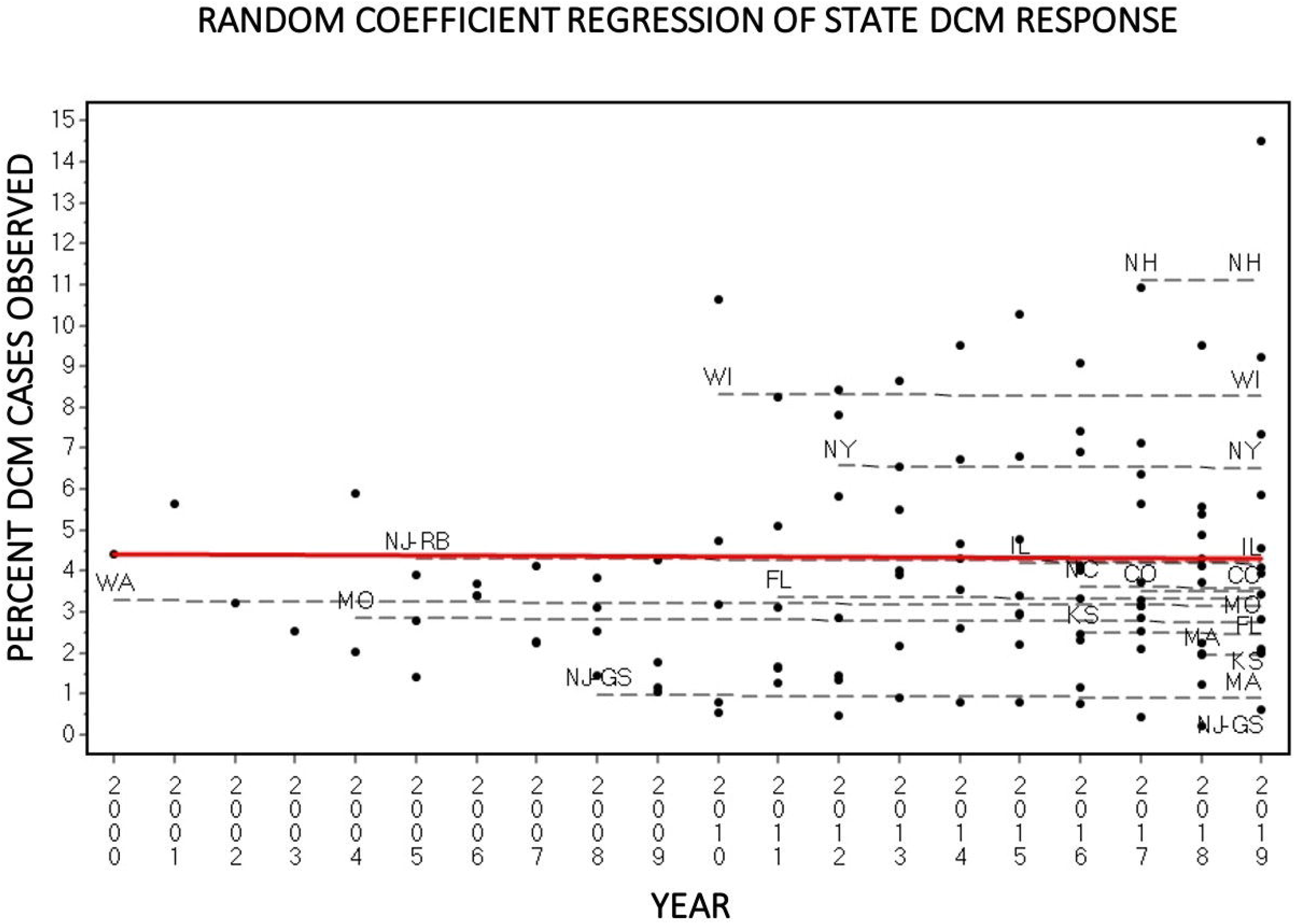
Linear regression model. Mixed model random coefficient regression analysis including all services with the exception of Ethos-Hawaii (only one data point). Black points indicate percent DCM per service each year. Trend - lines for each service are shown as black dashed lines. The best fitted regression model is shown in red. This trend line does not support a change in DCM incidence over time among cardiology services that participated in this study.

Cardiology services from Missouri and New Jersey (n = 2) provided additional data on annual breed predisposition (Fig 3). Fig 3A lists all breeds that were diagnosed with DCM phenotype from 2004-2019 from the three different cardiology services and represents how breeds were grouped for analysis. Breed distributions by site are displayed in scatter plots (Fig 3B-D). In all sites, breeds with a known inherited predisposition for DCM comprised the majority of cases diagnosed with DCM annually. Other breeds, including retrievers, mixed breeds, and to a lesser extent, small breeds, were also represented with lower frequencies over the past 15 years. A grouped Poisson regression analysis revealed positive trends for the Small Breed group and Other Breed group over the past 15 years, however these trends were not statistically significant (Small Breed, p=0.055, Other Breeds, p=0.053). A statistically significant positive trend was discovered in the Mixed Breed group (p=0.025), indicating that this group of dogs had an increased trend of DCM diagnoses over the past 15 years. There were no significant trends in the Inherited or Retriever groups (Fig 3D).

**Figure 3.**
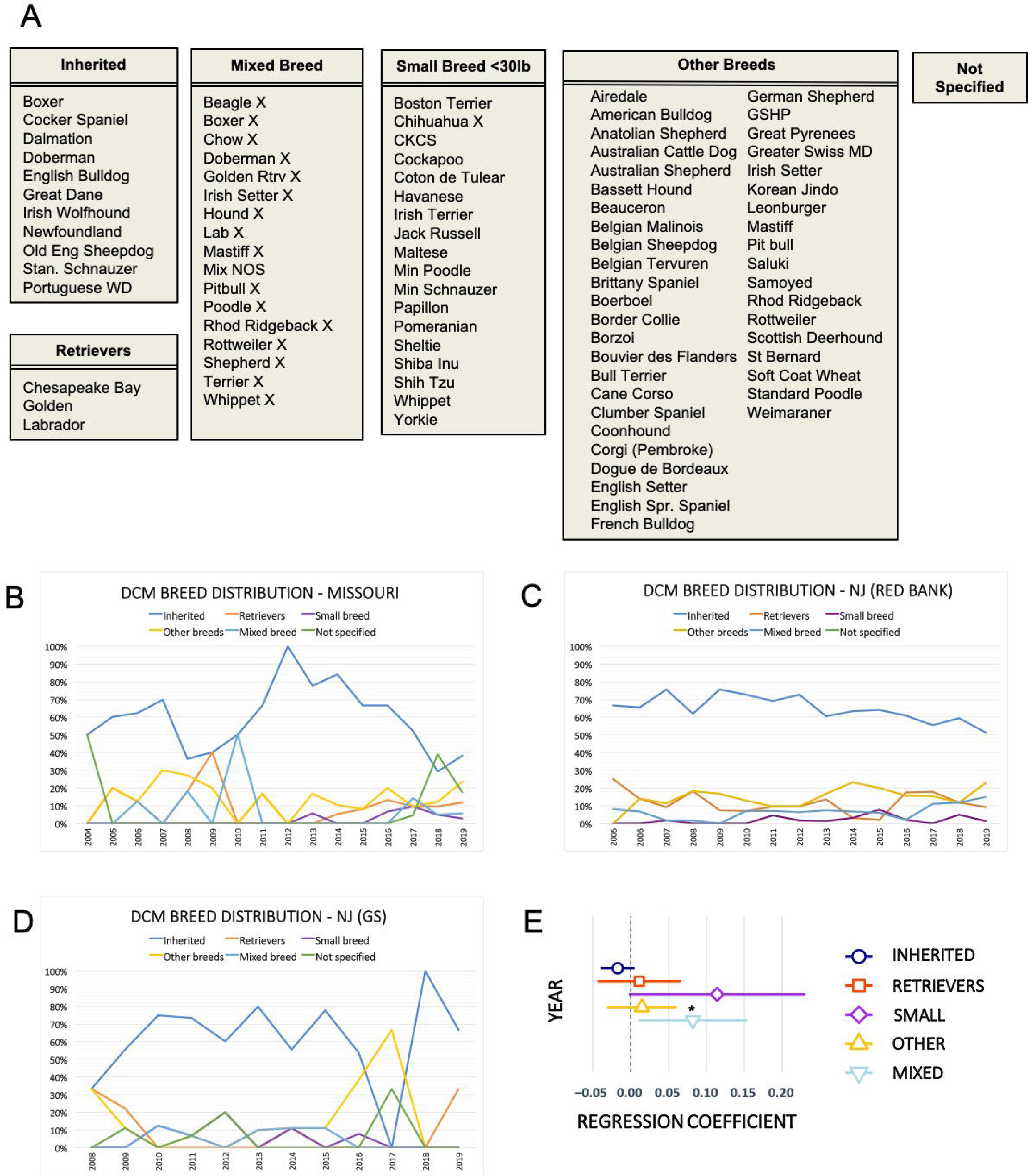
DCM breed distribution. Panel A. Dogs diagnosed with DCM were divided into 6 groups: 1. Inherited, 2. Mixed Breed, 3. Small Breed <30lb (13.6kg), 4. Other Breeds, 5. Retrievers, and 6. Not Specified. Panels B-D Annual breed distribution scatter plots from 3 different cardiology services: University of Missouri (B), Red Bank – New Jersey (C), and Garden State (GS) – New Jersey (D). Breeds with a known genetic link to DCM (Inherited group – dark blue) are the most commonly diagnosed with DCM, with all other categories generally being represented less frequently each year, in all three hospitals. Panel E depicts the results of a Poisson linear regression. The number of dogs in each group was regressed against year, total DCM cases, and site. Open icon denotes regression coefficient, with lines representing individual confidence intervals. Mixed breeds (light blue) showed a slightly increased trend, which was statistically significant (p=0.025, RC 0.082). Small breeds (purple) and other breeds (yellow) showed a statistical trend that was not significant (small breeds: p=0.055, RC 0.114; other breeds: p=0.053, RC 0.015). Retrievers (red) did not show a trend over the years (p=0.689, RC 0.0112). Inherited breeds (dark blue)showed a slight negative trend, which was not significant (p=0.134, RC - 0.017). RC= Regression Coefficient.

One service additionally provided annual age distribution (Red Bank - Tinton Falls, NJ) among DCM patients upon initial diagnosis, compared to all cardiology patients, and all patients annually evaluated by all services at the hospital (Fig 4A-B). An overall upward trend in age was noted among the three groups represented: All Cardiology Patients, DCM Patients, and All Canine Patients, over the past 15 years. DCM patients were consistently younger than other canine cardiology patients, and older than the general canine hospital population at the time of diagnosis. This trend was stable over time, though age of diagnosis increased within each of the three groups over time (Fig 4A). The age of DCM patients was divided into two groups: 0-6 years, and ≥7 years, and the trend was plotted over time (Fig 4B). In agreement with Fig 4A, there was a general upward trend in age at diagnosis of DCM in patients over the last 15 years. Whereas in 2005, 33% of DCM diagnoses were made at 0-6 years of age, and 67% of DCM diagnoses were made at ≥7 years of age, the age gap increased in 2020, with 25% diagnosed between 0-6 years of age, and 75% diagnosed at 7 or older, suggesting that dogs with DCM are living longer before diagnosis.

**Figure 4.**
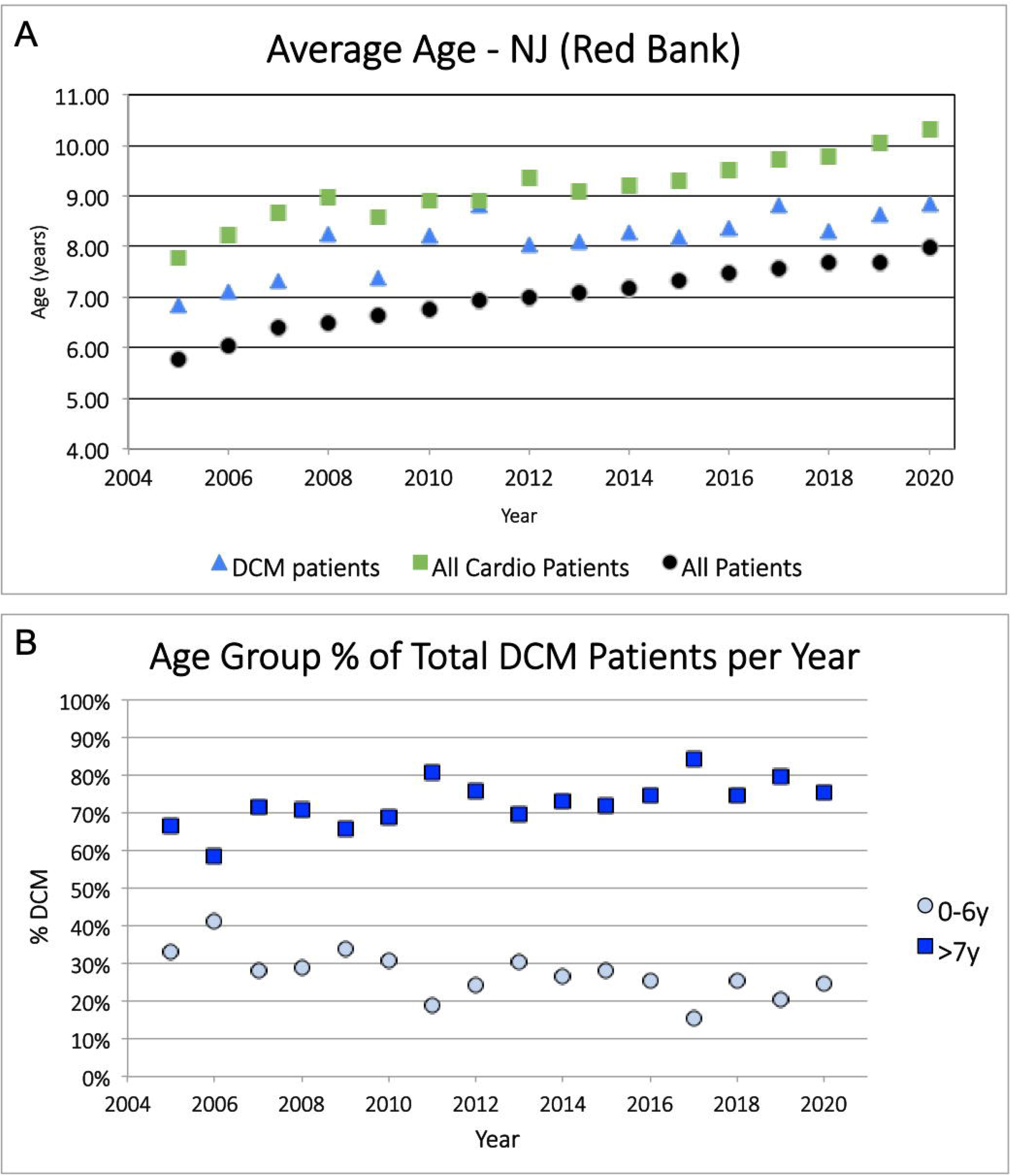
DCM age distribution. Age distribution of patients from Red Bank Veterinary Hospital (Tinton Falls) from 2005-2020. Panel A depicts the average age of patients at time of DCM diagnosis (blue triangles), at time of other cardiovascular diagnosis (green squares), or at the time of presentation for non-cardiac services (black circles). Within all three groups, there is an upward trend in age over the past 15 years. In general, patients diagnosed with DCM are one year older than the general population of canines presenting to the hospital, and one year younger than patients with an alternative cardiac diagnosis. Panel B depicts the percentage of DCM patients that are 6 years or younger (0-6y, grey circles), or 7 years or older (>7y, blue squares) at the time of diagnosis, from 2005-2020. The majority of patients (60% or more) are diagnosed with DCM at age 7 or above. In 2005, 67% of DCM patients were 7 years or older, while 33%were diagnosed at 6 years or younger. In contrast at year end 2019, 80 % of DCM patients were 7 years or older, and 20% were 6 years or younger. These data depict an increased age at initial DCM diagnosis over the past 15 years.

### Annual grain-free pet food sales

Data regarding grain-free pet food sales are fragmented, as markets that are not publicly traded are not required to report this information. Furthermore, the percentage of dogs eating grain-free diets is unknown. We obtained two data sets. First, the growth in annual grain-free pet food sales was provided by The Nielsen Company for the years 2011-2019 [11] (Fig 5A). These data included grain-free pet food sales from pet specialty suppliers only from 2011-2015, when sales increased from $900 million to $2.7 billion. Beginning in 2016-2019, the sales data additionally reflect brick and mortar food, drug, mass, and convenience stores (along with pet specialty sales), in accordance with the increasing popularity of grain-free diets during this time. Grain-free pet food sales increased to $5,426,879,686 in 2019. Importantly, not included in these sales data are grain-free pet foods sold by veterinary clinics, farm and feed stores, other stores, or online sales, likely resulting in an underestimate in total grain-free pet food sales from 2011-2019. Additional data from the Nielsen Company, xAOC, and Petco & Petsmart [11–13] providing percent market share of total grain-free pet foods, and grain-free dry dog food were evaluated from 2016 – 2019 (Fig 5B). Total grain-free pet food increased from 21.5% to 29% of the market share from 2016 to 2019, while grain-free dry dog food (the most common dog food type) increased from 31.9% - 43% of all kibble sold during this time (Fig 5B). Using the available data, we estimated that the percentage of US dogs eating grain-free diets was at least 2% in 2011 (based on the reported dollar sales of pet-specialty stores ($900 million), and the number of dogs in the US (78 million) [14]. In 2019, considering dollar sales alone ($5.5 billion) and the total number of dogs (90 million) [14], the percentage of dogs eating grain-free diets was estimated to be approximately 10%. However, when the percent market share of grain-free dry dog food was taken into account (43% in 2019), we estimated that the percentage of dogs eating grain-free diets could be as high 40% in 2019. While a five-fold increase in grain-free pet food sales was noted from 2011-2019, and with this, an increase in the percentage of dogs eating grain-free dog food is estimated, our data do not support a concurrent increase in cases of DCM (Fig 5C). Taken as a whole, these data do not support a correlation between grain-free diet sales and DCM incidence as diagnosed by veterinary cardiologists.

**Figure 5.**
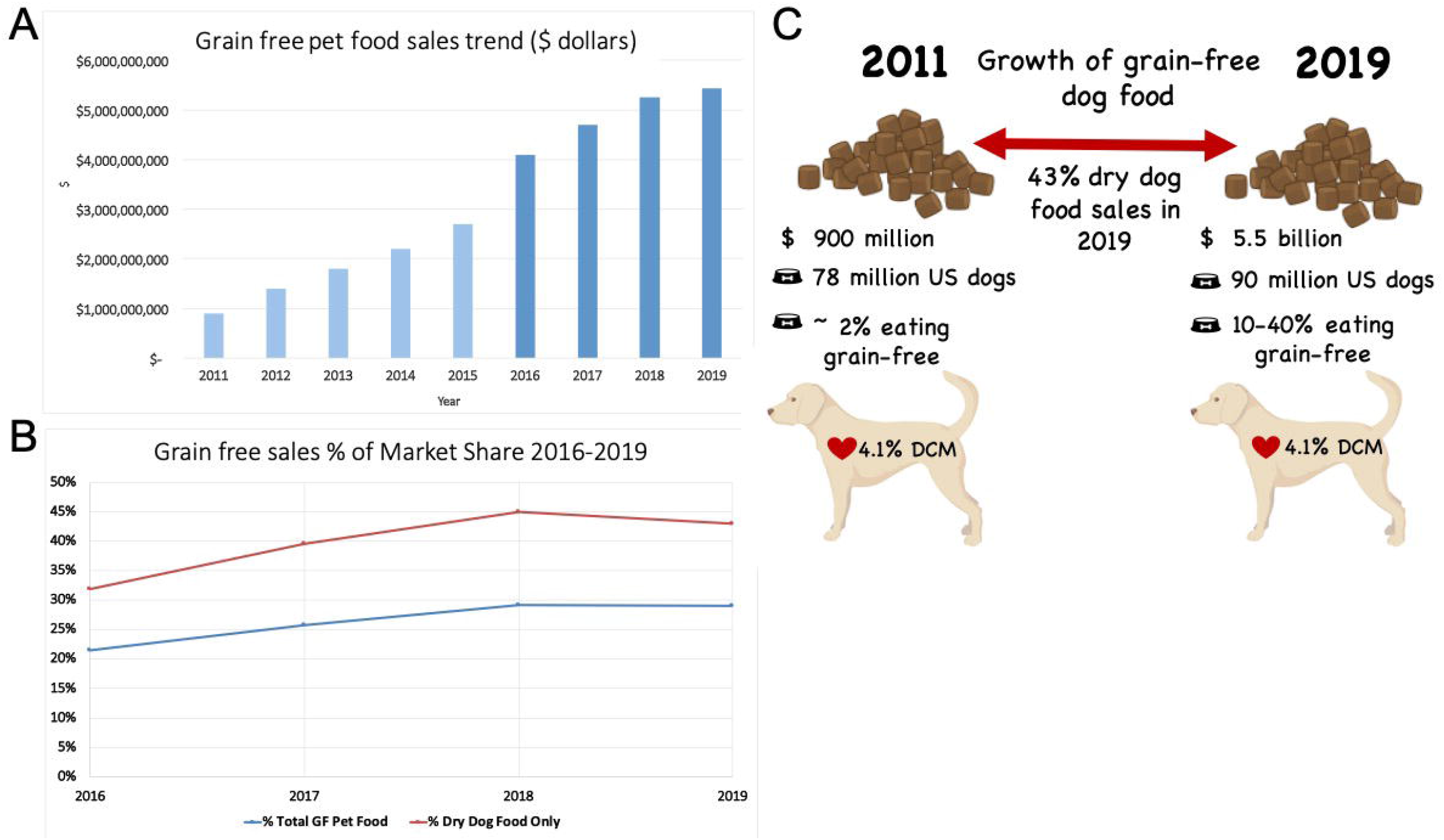
Pet food sales data. Grain-free pet food sales data provided by the Nielsen company, xAOC, and Petco & Petsmart, and used with permission. Panel A denotes grain-free pet food sales from 2011-2019. Grain-free pet food sales reached $900,000,000 in 2011, the first year that grain-free pet food sales were recorded. By 2019, sales had grown to $5,426,879,686. Light blue bars denote grain-free pet food sales through pet specialty retail stores only (2011-2015). Dark blue bars denote sales through brick and mortar food, drug, mass, and convenience stores *in addition to* pet specialty stores. These numbers are an underestimate of grain-free pet food sales, as sales through farm and feed stores, veterinary clinics, other stores selling pet foods, and online sales are not included. Panel B shows percent market share of grain-free pet food sales from 2016-2019. The blue line includes total grain-free pet food as a percent of the total pet food market. The red line denotes the percent of kibble that is grain-free from 2016-2019. Panel C shows a model, which estimates the percentage of dogs eating grain-free diets in 2011, compared to 2019. The percentage of dogs eating grain-free diets in 2011 was calculating using the following formula: $**900,000,000 grain-free sales / $50.00 per bag / 12 months / 78,000,000 dogs = 2%**. The percentage of dogs eating grain-free diets in 2019 was calculated using the same formula, adjusted for the number of dogs and dollar sales in 2019: $**5,5000,000,000 grain-free sales / $50.00 per bag / 12 months / 90,000,000 dogs = 10%.** These numbers are likely underestimates, as dollar sales do not include all sales of grain-free diets. Additionally, when considering the percent market share of grain-free dry dog food in 2019 (43%), the number of dogs eating grain-free kibble may be as high as 40% of US dogs. In contrast, the average % DCM diagnosed among cardiologists remained stable at 4.1% during this time period.

## Discussion

Here, we provide a retrospective data set that describes the overall incidence of DCM as diagnosed by 14 veterinary cardiology services across the US over the past two decades. Variation in DCM incidence rates is present at different cardiology services. This is expected, due to multiple regional variables, not limited to: variation in diagnostic methods among cardiologists, different breed distributions across states, regional infectious diseases that may result in myocarditis and other systemic disease, and pet owner demographics. Despite the state-by-state variation in annual DCM incidence, the overall trend does not support a change in DCM incidence, with the average annual incidence rate at 3.83% (Range 2.41-5.65%, STD +/- 0.77%) over the past 19 years. This average is considerably higher than what has been reported previously [2,10], however, our data reflects DCM diagnosed as a proportion of cardiology cases only, rather than DCM within the whole population. Interestingly, data from one hospital that provided total hospital caseload numbers from 2005-2020 revealed a similar incidence rate to what has been previously reported (Fig 1B), suggesting that the DCM incident rate may be on trend today to what was documented from 1986-1991 [2] and 1995-2010 [10]. Detailed statistical analysis revealed that a group of cardiology services (FL, KS, NC, NJ (2), NY, WI) reported a statistically significant decrease DCM, while other services (CO, IL, MA, MO, NH, WA) reported an upward (but not significant) trend. While these data are interesting, they are difficult to interpret as a particular pattern is not evident to explain these trends. The states, as grouped, do not comprise particular geographical areas of the US, nor is there a clear relationship between them. One explanation for these varying trends is the small data set that is being analyzed. It is possible that if a longer time range were available for analysis among a larger number of services, that the trends would become clearer. Additionally, individual breed popularity by state and regional infectious disease prevalence, as well as other unknown variables, may also play a role in these trends but were not evaluated in this study.

Additionally, we provide data describing the breed distribution of dogs diagnosed with DCM across three hospitals. Evaluation of the available breed predisposition data supports previous studies that describe DCM as a primarily inherited disease, with breeds with a known inherited predisposition comprising the majority of diagnosed DCM each year, in each cardiology practice. Furthermore, retrievers, mixed breed dogs, other large breed dogs, and small breed dogs have historically been diagnosed with DCM, to a lesser extent, for the past 15 years. However, a regression analysis revealed that only the Mixed Breed group had a statistically significant upward trend. Positive trends that were not statistically significant were noted in the Small Breed and Other Breed groups. These data may support what has been reported by some cardiologists as an increase in atypical breeds developing DCM phenotype [7–9, 15, 16]; however, these data sets are few, and do not support a causative relationship between DCM phenotype and any specific variables. It should also be noted that some dogs in the Mixed Breed group were noted to be mixed with a breed with a known predisposition to DCM (Fig 3), and it is possible that an increased popularity of mixed breed dogs over the past two decades [17,18] has contributed to this upward trend. Additionally, we characterized dogs in the Inherited group as those with a published record of inherited disease. Breeds, such as the Scottish Deerhound, St. Bernard, and Saluki have been reported to be overrepresented [19], however, were placed into the Other Breeds category, as not enough data are available at this time to confirm a hereditary predisposition.

One veterinary hospital provided age distribution date over the past 15 years (2005-2020). These data provide fascinating insight into the age at which dogs are diagnosed with cardiac disease as a whole, compared to the age of the general population of dogs seen by all services at the hospital. In general, across all categories, there is an upward trend to the age at which dogs are evaluated at a specialty and emergency center over the past 15 years, or simply put, our dogs are living healthier for longer. This likely results from several factors including: 1) advancements in medical therapy, 2) increased availability of veterinary care, and 3) increased desire and financial capability of owners to seek advanced medical care for their pets. Compared to the general population, dogs diagnosed with DCM are approximately one year older – this trend continues over 15 years. In contrast, compared to the population of dogs diagnosed with other cardiovascular disease, dogs diagnosed with DCM are approximately one year younger. This is in agreement with past studies that describe the average age of onset of DCM being 6-8 years of age [20, 21], while the most common cardiac disease diagnosed in the dog, degenerative mitral valve disease [22], tends to have an average onset of 8-10 years of age [22]. It is interesting to consider that, despite the dramatic increase in grain-free pet food sales over the past decade, changes in age distribution over time were not observed. Assuming that dogs of all ages are fed grain-free diets, and that it has been hypothesized that it may take a few as three months to develop nutritional cardiomyopathy secondary to these diets [23], one might expect that the average age of diagnosis of DCM might decrease over time (as younger dogs developed the disease) - if a large enough percentage of dogs were eating grain-free diets and a true correlation existed. Such a trend was not observed with the data, though the actual percentage of dogs eating grain-free diets at this practice during this time period is unknown.

Concern over the relationship between certain diets and DCM has risen over the past few years [7–9, 24–28]. An increase in total DCM cases in Golden retrievers eating grain-free diets has been reported in peer-reviewed literature [7,16,23], though recently an Expression of Concern was reported by the editors [29], leading to further confusion regarding the issue of diet and DCM. Previous studies have also suggested a genetic correlation to taurine deficiency in the golden retriever, irrespective of diet [16, 30]. While our data reveals that retrievers (including goldens, Labradors, and Chesapeake Bay) have been diagnosed with DCM over the past 15 years, an increased incidence was not supported. Further anecdotal reports state cases of DCM increasing in atypical breeds [7–9,24–27], but no measurable data was previously available to support these claims. Our data support a weak upward trend in “atypical breeds,” including mixed breeds (statistically significant trend), and other small and large breeds (not statistically significant) developing DCM over the past 15 years. In addition to external factors such as changes in breed popularity, infectious disease, and diet, it must be considered that an increase in atypical breeds may simply be a reflection of previously unrecognized heritable cardiomyopathies occurring in the breeds as seen with toy Manchester terriers, standard schnauzers, Welsh springer spaniels, Rhodesian ridgebacks, Dalmatians, and English bulldogs [31–37]

Here, we describe a more than five-fold increase in grain-free pet food sales over the past decade, with no correlating increase in overall DCM incidence. The number of dogs currently eating grain-free diets is unknown, but we estimate it to be between 10-40% of the US dog population (Fig 5C). The massive increase in grain-free diet sales over the past decade means it increasingly likely that dogs will be consuming these foods. Furthermore, grain-free diets are generally considered premium dog foods and are often more expensive than their grain-inclusive counterparts. Therefore, dogs seen by veterinary specialists may be even more likely to be consuming grain-free diets, as they represent a different demographic than the general population. However, an increased likelihood of eating grain-free dog food does not necessarily correlate to increased risk of DCM, and a causative link between grain free diets and DCM has not yet been shown. While anecdotal reports of dogs improving after diet change have been reported as evidence that specific diets were the cause of DCM phenotype [7–9,16, 38] close evaluation of the data reveals multiple variables at play, including initiation of taurine and carnitine supplementation, and medications to treat congestive heart failure and tachyarrhythmias [7–9, 16, 28, 38, 39]. All of these factors may play a role in the improvement of cardiac function in cases of DCM phenotype. Furthermore, spontaneous resolution of DCM phenotype had been reported in cases of viral myocarditis, in both humans [40] and dogs [41, 42]. These factors must all be considered when trying to determine the underlying primary cause for DCM phenotype.

The authors understand there are many limitations when conducting a retrospective study based on referral case numbers of multiple independent institutions. Some of those include collecting sufficient information from every region of the country, collecting sufficient numbers of years of data from each institution to understand the trend of DCM diagnoses for that particular hospital, and varying criteria for the diagnosis of DCM for each individual referral hospital. More specifically, despite our best efforts, we were able to obtain data from fewer than 10% of board-certified veterinary cardiologists in the US, and while this is the most complete dataset available at this time, it is possible that it does not represent the trends observed by all veterinary cardiologists. The small numbers of animals considered in the breed distribution data are interesting to consider; however, may not give an accurate picture of breed distribution at all cardiology services. Furthermore, the data do not consider risk factors for small cohorts of breeds, which may be excluded from this global study due to small numbers. For example, the golden retriever is overrepresented in all datasets in all years, compared to all other breeds. The authors suggest that this may indicate an, as-of-yet undetermined (though previously suspected) hereditary component to DCM in this particular breed and that golden retrievers may have different dietary requirements regardless of diet [16, 30]. Additionally, the presence of a hereditary component in any of the “atypical breeds” (including mixed breeds) that appear to be overrepresented for DCM should be considered. Finally, the presence of other well-established causes of DCM phenotype, such as myocarditis, hypothyroid disease, or chronic tachycardias should not be ignored when considering the etiology of DCM phenotype. Further data from veterinary cardiologists including all cardiology patients and their detailed diet histories would allow us to assign hazards ratios to these variables.

Limitations notwithstanding, this data comprises the most current data set considering the incidence, age, and breed distributions of canine DCM diagnosed by veterinary cardiologists.

---

**Supplemental Figure 1.**

DCM Incidence Survey provided to veterinary cardiologists.

**Supplemental Table 1.**

Linear regression data for individual services, all services, and services grouped by trend. n= number of years for which data was provided. Slope and p values are also provided. Statistically significant trends are denoted with an *. Individually, both Garden State (NJ), and SW Florida Vet Specialists had statistically significant negative trends. When sites were grouped with like negative or positive trends, the negative trends group (FL, KS, NC, NY, NJ-GS, NY-RB, WI) was statistically significant, but the positive trends group (CO, IL, MA, NH, WA) was not. Multiple testing corrections were not performed, but these trends are likely not robust enough to maintain significance. When grouped as a whole, there was no significant trend in % over time.

## Acknowledgements

We would like to thank the following individuals and institutions, without whose assistance, this project would not have been possible:

Dr. Justin Thomason DVM, DACVIM - Cardiology (Kansas State)

Dr. Wendy Arsenault DVM, DACVIM - Cardiology (SW Florida Veterinary Specialists)

Dr. Lynne Nelson DVM, DACVIM - Cardiology (Washington State)

Dr. Ryan Baumwart DVM, DACVIM - Cardiology (Washington State)

Dr. Samuel D. Stewart, DVM, DACVECC (Ethos Veterinary Health)

Dr. Lindsey Bullen DVM, DACVN (VSH North Carolina)

Dr. Elizabeth Lund DVM, MPH, PhD (Compassion -First Pet Hospitals)

Mary Semplenski, Report Analyst (Compassion -First Pet Hospitals)

Edwin Ortiz, Manager, Clinical Studies and Social Responsibility (Compassion -First Pet Hospitals)

Dr. Jonathan Goodwin DVM, DACVIM - Cardiology (Veterinary Specialty Network)

Dr. Max White DVM, DACVIM - Cardiology (Garden State Veterinary Specialists, Tinton Falls, NJ)

Kristie Garcia, LVT, VTS – Cardiology (Garden State Veterinary Specialists, Tinton Falls, NJ)

Finally, thank you to Dr. Jim Schwenke and Dr. Charles Danko for assisting with the statistical analysis.

