## Supplemental Figure 1 for "Incidence of canine dilated cardiomyopathy, breed and age distributions, and grain-free diet sales in the United States from 2000-2019: A retrospective survey"

BSM Research™
a BSM Partners Practice Group

### DCM INCIDENCE SURVEY

Goal: To identify the incidence of canine dilated cardiomyopathy diagnosed by veterinary cardiologists in the United States over the past 10-15 years.

Name of Hospital:

Number of board-certified cardiologists:

Number of residency-trained cardiologists:

Number of residents:

Criteria your group uses for diagnosing DCM (*please highlight any that are used* ***in the majority*** *of cases*):

FS% <

EF%<

LVIDDn >

LVIDSn >

Sphericity index

EPSS

Tissue Doppler

Strain/Strain rate

### DCM INCIDENCE SURVEY

Subjective assessment of the left ventricle

Breed

Other:

What operating system/database was used to collect your data?

Please describe the search criteria you used to amass this data: *(ex. search terminology such as dilated cardiomyopathy, DCM, breed, age, etc…)*

Have you included dogs with a description of “decreased systolic function” in this search, or did you focus on the search terminology “dilated cardiomyopathy” only?

Would it be possible to share additional data such as breed, age, and gender with us to expand our database?
