## Supplementary figures and images for "Incidence of canine dilated cardiomyopathy, breed and age distributions, and grain-free diet sales in the United States from 2000-2019: A retrospective survey"

### Supplemental Table 1

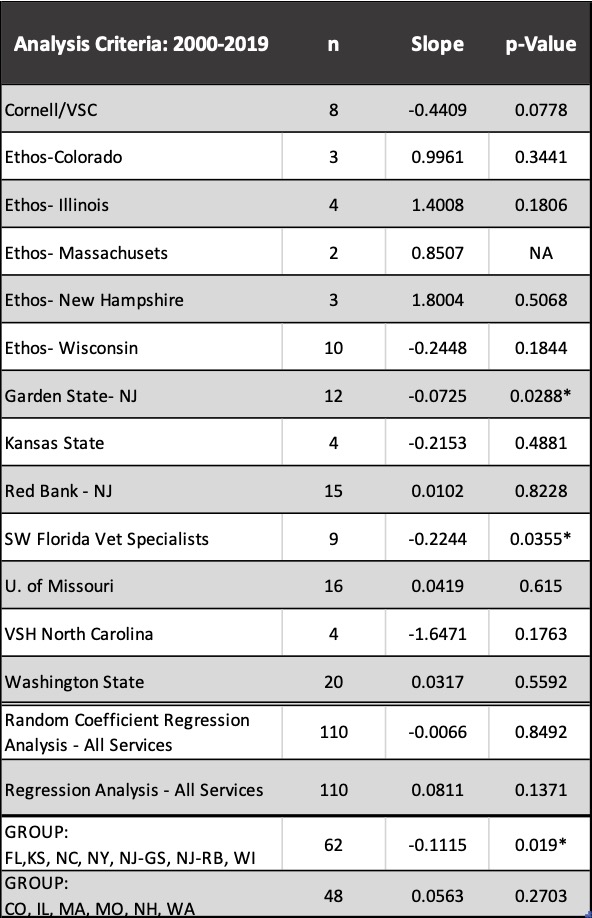
